# Methods for recording and imaging spreading depolarizations and seizure-like events in mouse hippocampal slices

**DOI:** 10.1101/2021.08.19.457007

**Authors:** Yi-Ling Lu, Helen E. Scharfman

## Abstract

Spreading depolarization (SD) is a sudden and synchronized depolarization of principal cells followed by depression of activity, which slowly propagates across brain regions like cortex or hippocampus. SD is considered to be mechanistically relevant to migraine, epilepsy, and traumatic brain injury. Interestingly, research into SD typically uses SD triggered immediately after a focal stimulus. Here we optimize an in vitro experimental model allowing us to record SD without focal stimulation. This method uses electrophysiological recordings and intrinsic optical imaging in slices. The method is also relatively easy and inexpensive. Acute hippocampal slices from mice or rats were prepared and used for extracellular and whole-cell recordings. Recordings were made in a submerged-style chamber with flow of artificial cerebrospinal fluid (aCSF) above and below the slices. Flow was fast (> 5ml/min), and temperature was 32°C. As soon as slices were placed in the chamber, aCSF containing 0 mM Mg^2+^ and 5 mM K^+^ (0 Mg^2+^/5 K^+^ aCSF) was used. Two major types of activity were observed: SD and seizure-like events (SLEs). Both occurred after many minutes of recording. Although both mouse and rat slices showed SLEs, only mouse slices developed SD and did so in the first hour of 0 Mg^2+^/5 K^+^ aCSF exposure. Intrinsic optical imaging showed that most SDs initiated in CA3 and could propagate into CA1 and dentate gyrus. In dentate gyrus, SD propagated in two separate waves: (1) into the hilus and (2) into granule cell and molecular layers simultaneously. This in vitro model can be used to better understand the mechanisms and relationship between SD and SLEs. It could also be useful in preclinical drug screening.

## 1 Introduction

Increased neuronal excitability is a common feature underlying several neurological disorders, including but not limited to migraine, epilepsy, traumatic brain injury, and stroke (Eikermann-Haerter et al., 2013; Noseda and Burstein, 2013; Rogawski, 2013; Kim et al., 2014; Bugay et al., 2020). Although seizures may be the best-known example that results from increased neuronal excitability, spreading depolarization (SD) is another example.

SD is a group of cells that depolarize toward 0 mV when ion gradients across membrane are almost completely lost (Somjen, 2001; Hartings et al., 2017). When electrical activity is monitored extracellularly, this dramatic depolarization features a significant negative potential shift in an extracellular recording (direct current (DC) recording). The DC negative shift is followed by an extended suppression of spontaneous electrical activity. Leão first reported the activity suppression after focal electrical stimulation and how it slowly spread across anesthetized rabbit cortex (Leão, 1944). He coined the term “spreading depression (SD).” However, after insults to human brain, the suppression of spontaneous activity can happen before the depolarization (Dreier, 2011; Ayata and Lauritzen, 2015). Below we use the term SD to refer to more than one type of SD.

As membrane potential depolarizes, neurons begin action potential (AP) firing which stops as the cell reaches its most depolarized potential. Neurons also show reduced input resistance (Czéh et al., 1993). There is extensive influx of sodium and calcium, and large efflux of potassium, leading to a temporary loss of ionic gradients across the membrane (Somjen, 2001; Dreier, 2011; Ayata and Lauritzen, 2015). The inflow of cations brings water inside the neuron, which leads to swelling and alters the light transmission through tissue (Dreier, 2011). Therefore, the change in light transmission has been used as a method to monitor SD-associated swelling and the propagation of SD from one brain area to the next (e.g., Anderson and Andrew, 2002; Buchheim et al., 2002). Remarkably, neurons often recover after glia and ion pumps restore normal ionic gradients (Lian and Stringer, 2004; Dreier, 2011; Varga et al., 2020).

To gain a better understanding of mechanisms underlying SD, past hippocampal studies have typically used slice models and rats. Many of these studies were conducted in an interface-style chamber (e.g., Snow et al., 1983; Scharfman, 1997; Buchheim et al., 2002; Reyes-Garcia et al., 2018). In an interface-style chamber, brain slices lie on a mesh at the interface between artificial cerebrospinal fluid (aCSF) and air. Warm, humidified O_2_ or 95% O_2_/5% CO_2_ is vented over the slices (Schwartzkroin, 1975; Haas et al., 1979; Scharfman et al., 2001). Although interface-style chambers are widely used to study SD, lack of a powerful microscope limits visualization (but see Case and Broberger, 2013). On the other hand, a submerged-style chamber that is typically used with a high-resolution microscope allows visualization of neurons. Visualization is an advantage because it facilitates patch clamp recordings and optical imaging.

In both types of recording chambers, SDs are often initiated by increasing local excitability using electrical stimulation or focal high concentration of potassium (e.g., Czéh et al., 1993; Buchheim et al., 2002; Lindquist and Shuttleworth, 2012; Eickhoff et al., 2014; Martens-Mantai et al., 2014). When a different approach is used to increase excitability, such as reduced magnesium in aCSF, a focal stimulation can elicit an SD when slices are in an interface-style chambers (Scharfman, 1997) but only a SLE when a submerged-style chamber is used (Anderson et al., 1986; Lewis et al., 1990; Churn et al., 1991; Kojima et al., 1991). Delayed, spontaneously developed SD using a low magnesium aCSF is rarely reported. When reported, an interface-style chamber was used (Mody et al., 1987). Therefore, methods using a submerged-style chamber to examine SD with altered aCSF would be advantageous.

Here, we report an optimized method to examine SD that occurs without a time-locked focal stimulus, which we refer to as “spontaneous” SD for simplicity. These spontaneous SDs appeared to be spontaneous because they had a long and unpredictable delay before SD ultimately occurred. We prepared slices in sucrose-containing aCSF (see Methods). When they were placed in the recording chamber, we began to use aCSF containing nominal Mg^2+^ and higher K^+^ (0 mM Mg^2+^, 5 mM; 0 Mg^2+^/5 K^+^ aCSF). A previous report has shown that 0 Mg^2+^/5 K^+^ aCSF can slowly change the slice in an interface-style chamber so that eventually SD and SLEs develop, and a focal stimulation is optional (Mody et al., 1987). Our methods allowed us to record both spontaneous SDs and SLEs, which is valuable because the interplay between them is rarely studied, and therefore is poorly understood. We added intrinsic optical imaging and developed methods to quantify the imaging results as well as simultaneous recordings of SD and SLEs. These methods provided additional insight about SD beyond what is known. We suggest that these methods can help provide further advances in understanding SD as well as providing a preparation that may be useful in preclinical drug screening.

## 2 Materials and Methods

The experimental procedures were carried out in accordance with the National Institutes of Health guidelines and were approved by the Institutional Animal Care and Use Committee in The Nathan Kline Institute. All chemicals were obtained from Millipore-Sigma unless otherwise specified.

### 2.1 Animals and husbandry

A total of 10 Sprague-Dawley rats (all males, age 21 – 45 days) and 26 C57BL/6 mice (14 males and 12 females, age 20 – 36 days) were used in the current study (Table 1). The maximum number of animals housed per cage was three for rats and four for mice. All animals were housed using a 12 hr light/dark cycle. Food (Purina 5001 Chow, W.F. Fisher) and water were available ad libitum. Numbers of animals that were used in each experimental condition are listed in the Table 1.

**Table 1.**
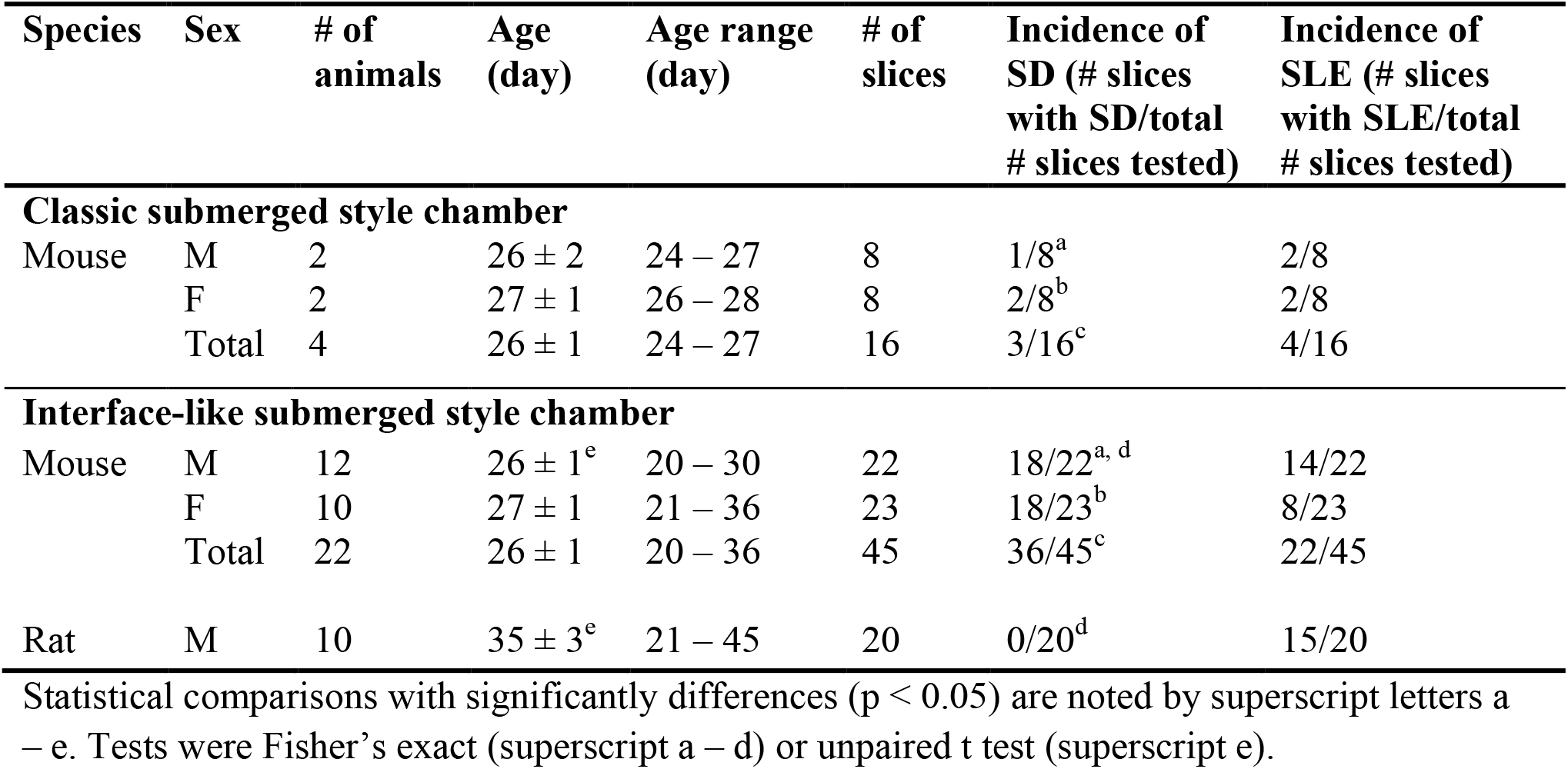
Conditions tested for the development of SD and SLE using 0 Mg^2+^/5 K^+^ aCSF.

### 2.2 Terminology

Definitions of the terms described below were defined based on past studies of SD and our data. Extracellularly, **SD** was defined as an event that consists of a negative DC shift with a slow recovery. In our model, SD almost always began with a series of high frequency bursts and then proceeded to a negative DC shift. Intracellularly, SD began with a short period of depolarization over 20 mV with high frequency firing of APs. Subsequently, there was a slow repolarization.

A **SLE** was defined by multiple criteria to distinguish it as abnormal and seizure-like. Extracellularly, SLEs were defined as multiple bursts of population spikes superimposed on short-lasting positive waves. In slices, where spontaneous population spikes are not normally present, SLEs were easy to distinguish. SLEs were referred to as seizure-like because they showed more and longer trains of bursts than the 1 – 2 bursts referred to as an epileptiform discharge. SLEs also were composed of bursts at high frequency (> 1 Hz) and lasted much longer than epileptiform burst discharges (Borck and Jefferys, 1999). Intracellularly, the bursts of an SLE corresponded to an initial depolarization with APs at the peak, somewhat like a paroxysmal depolarization shift (PDS) (Borck and Jefferys, 1999). The SLEs observed in our preparation resembled events that have been called ictaform events or ictal activity/discharge in the past (Anderson et al., 1986; Avoli et al., 2002).

### 2.3 Slice preparation and electrophysiological recording

Rats or mice were deeply anesthetized using isoflurane (Patterson Veterinary) and then decapitated. Brains were quickly removed and immersed in ice-cold sucrose-containing aCSF (sucrose aCSF, ingredient in mM: 90 sucrose, 80 NaCl, 2.5 KCl, 1.25 NaH_2_PO_4_, 25 NaHCO_3_, 10 D-glucose, 4.5 MgSO_4_, and 0.5 CaCl_2_, pH = 7.3 – 7.4). Horizontal hippocampal slices (350 µm thick) were obtained using an oscillating tissue slicer (Microm, HM650V, Thermo Fisher Scientific, or VT1200 S, Leica). Brain slices were then transferred to a holding chamber (made in-house) containing sucrose aCSF. This chamber allowed slices to sit on a mesh several inches from the base of the chamber but still below the surface of aCSF, and the aCSF circulated around the slices. The holding chamber with brain slices was placed in a water bath and temperature was gradually increased to 35°C. Afterwards, the temperature of the water bath was maintained at 35°C for 45 min (“recovery”). After recovery, slices were maintained at room temperature in sucrose aCSF for the rest of the day (Figure 1A). All aCSF for slicing and recording was constantly oxygenated using carbogen (95% O_2_ and 5% CO_2_).

**Figure 1.**
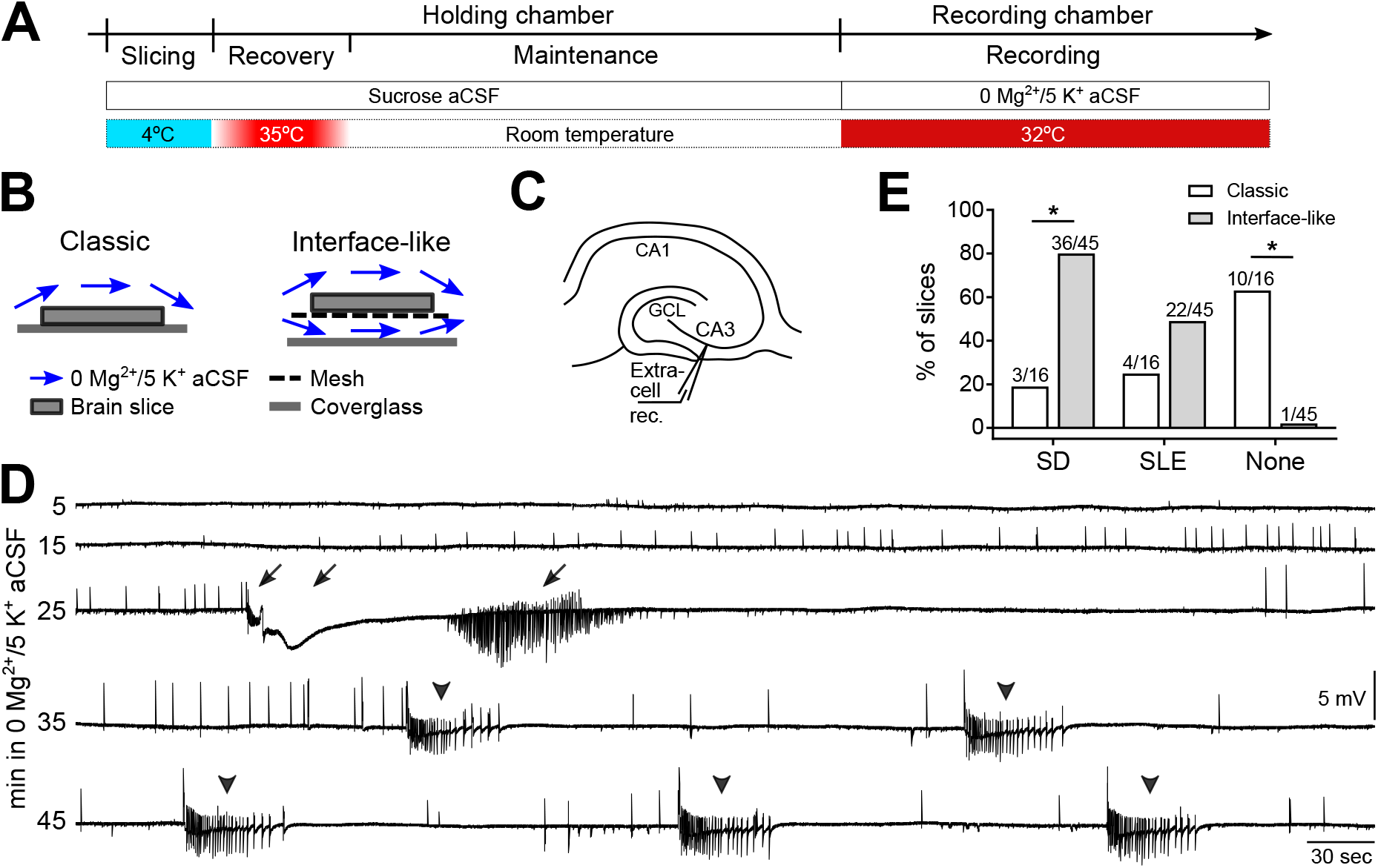
SD and SLEs are developed in 0 Mg^2+^/5 K^+^ aCSF in an interface-like submerged-style chamber. **(A)** Experimental timeline. Hippocampal slices were prepared in ice-cold sucrose-based artificial cerebrospinal fluid (sucrose aCSF). Slices were incubated at 35°C for 45 min and then maintained at room temperature in sucrose aCSF before being transferred to the recording chambers for recording at 32°C. **(B)** Illustration of classic (single superfusion) and interface-like (dual superfusion) recording chambers. **(C)** Schematic of extracellular recording in the CA3 of hippocampus. **(D)** A representative trace shows a continuous recording of extracellular activity in the CA3 of hippocampus in 0 mM [Mg^2+^]_o_ /5 mM [K^+^]_o_ aCSF (0 Mg^2+^/5 K^+^ aCSF). The slice was recorded in an interface-like recording chamber. Both spreading depolarization (SD) and seizure-like event (SLE) developed, and either one could precede the other. Arrows point to a single SD. Arrowheads point to five SLEs. **(E)** Percentage of SDs, SLEs, and neither event (None) developed in classic (white bars) or interface-like (gray bars) recording chambers within the first hour of 0 Mg^2+^/5 K^+^ aCSF exposure. Slices recorded in an interface-like recording chamber showed a higher percentage of SD than in a classic recording chamber (Two-sided Fisher’s exact test, * p < 0.0001). Nighty-eight percent of slices recorded in an interface-like chamber developed SD and/or SLE.

Brain slices were transferred from the holding chamber to a recording chamber for data collection. Two types of recording chambers were used in the current study: a “classic” submerged-style chamber (classic chamber, RC-26GLP, Harvard Apparatus, Figure 1B) and an “interface-like” submerged-style chamber (interface-like chamber, RC-27LD, Harvard Apparatus, Figure 1B). When slices were placed in the recording chamber, we used aCSF with 0 mM MgSO_4_ and 5 mM KCl (0 Mg^2+^/5 K^+^ aCSF) as follows (in mM): 130 NaCl, 5 KCl, 1.25 NaH_2_PO_4_, 25 NaHCO_3_, 10 D-glucose, 0 MgSO_4_, and 2.4 CaCl_2_. Recordings were performed at 32 ± 1°C with 5 – 7 mL/min flow rate.

Recording electrodes were pulled (P-97, Sutter Instruments) from borosilicate glass (1.5 mm outer diameter, 0.86 mm inner diameter, Sutter Instruments). An extracellular recording electrode containing 0 Mg^2+^/5 K^+^ aCSF (resistance 3 – 7 MΩ) was placed in CA3 stratum pyramidale to record field potentials. For two simultaneous extracellular recordings, a second recording electrode was placed in either CA1 or the granule cell layer (GCL) at the crest of the dentate gyrus. In some slices, a second recording electrode (resistance 5 – 8 MΩ) was used to make whole-cell recordings of individual CA3 pyramidal cells simultaneously. The internal solution for whole-cell recording contained (in mM) 120 K-gluconate, 10 HEPES, 20 KCl, 2 MgCl_2_, 0.2 EGTA, 4 Mg-ATP, 0.3 Na_2_-GTP, 7 Tris-phosphocreatine, and 0.5% biocytin. Data were amplified (MultiClamp 700B, Molecular Devices), digitized at 10 kHz (Digidata1440A or 1550B, Molecular Devices), and acquired using pClamp (v. 10.7 or 11, Molecular Devices).

### 2.4 Biocytin labeling

Immediately after recording, hippocampal slices were rinsed in 0.9% saline (all solutions used deionized water; dH_2_O) and then preserved in 4% paraformaldehyde in 0.1M phosphate buffer (pH 7.4). Slices were stored at 4°C until they were processed. Procedures were conducted using free-floating sections at room temperature with mild agitation using a clinical rotator (The Belly Dancer, Stovall Life Science Inc.) unless otherwise specified. For the following steps, all buffers were prepared in 0.1 M Tris buffer (TB, pH 7.6). First, slices were incubated in 1% Triton-X 100 (Triton) for an hour to make cell membranes more permeable to reagents. Then endogenous peroxidase activity was suppressed by incubating slices in 0.1% H_2_O_2_ for 30 min. After three rinses (10 min each) with 0.25% Triton, slices were incubated in an avidin-biotin complex solution (ABC Elite Kit, Vector Laboratories) overnight at 4°C with gentle rotation. Slices were pretreated with 0.05% 3,3’-diaminobenzidine (DAB) with 1 mM NiCl_2_ for 30 min followed by incubation of 0.05% DAB with 0.0075% H_2_O_2_ (diluted in dH_2_O) until biocytin-filled cells were clearly visualized. The DAB reaction was terminated by three 10 min washes in TB. Slices were equilibrated with increasing concentrations of glycerol (15 min each, 25%, 40%, 55%, 70%, 85%, 100%) (Hamam and Kennedy, 2003). Slices were coverslipped in 100% glycerol and edges of coverslip were sealed using nail polish. Brightfield images were taken using a charge-coupled device (CCD) camera (Retiga R2000, Teledyne QImaging) and an upright microscope (BX61, Olympus) with a 10X objective (UPlanSApo, 0.40 N.A., Olympus). ImagePro 7 Plus (Media Cybernetics) software was used for image acquisition. Brightfield images from different focal plans were stacked using CombineZP software (Hadley, 2010; Botterill et al., 2017) or the Stack Focuser plug-in in ImageJ (Umorin, 2002; Schneider et al., 2012).

### 2.5 Intrinsic optical imaging

Spontaneous SD and SLEs in CA3 were continuously monitored using field potential recordings. Recording of intrinsic optical signals was started manually as soon as electrophysiological manifestations of SD were observed. The time of beginning of the imaging was recorded and matched with electrophysiological recordings. One slice was excluded from imaging analysis because the imaging was started after the beginning of SD.

Intrinsic optical signals were captured using time-lapse images at 1 or 2 frames per second. All images covered each of the major subfields in hippocampus (dentate gyrus, CA3, CA2, CA1, subiculum) and some of the cortex. The first image of an event was used as a baseline for comparisons of changes in intrinsic optical signals within the event. This method to define a baseline was chosen because the first three images showed minimal or no change in intrinsic optical signals.

Time-lapse images were acquired every 0.5 s for eleven slices and 1 s for two slices. The overall propagation patterns of imaged SDs were similar whether acquisition was 0.5 or 1 sec/image, so the data using the two different acquisition rates were pooled. A CCD camera (Retiga Electro, Teledyne Photometrics) mounted on an upright microscope (BX51, Olympus) with a 4X objective was used for image acquisition. No multiplier was used between objective and the camera. A 775 nm filter was used to allow red/far red light to pass through the recorded slice.

The light intensity was determined at the time when a slice was transferred into the recording chamber without saturated pixels. The same light intensity was used throughout the recording of a given event. Exposure time of each image was 10 msec. MicroManager (Edelstein et al., 2010, 2014) or Ocular (Teledyne Photometrics) was used for image acquisition. Images were stored in stacks and analyzed using ImageJ or Fiji (Schindelin et al., 2012; Schneider et al., 2012).

To visualize changes in light transmission in pseudocolor, several steps were taken. First, images during an event (SD or SLE) were subtracted from the baseline image. Specifically, Image Calculator in ImageJ was used to calculate the difference between an image stack of an event (Event Stack) and an image stack generated from the event’s baseline image (Baseline Stack).

The result was a stack of images that showed the difference from baseline in light intensity (Result Stack). Next, images from Result Stack were pseudocolored for better visualization where warmer and cooler colors indicated higher and lower light transmission, respectively. The Event Stack was later combined with its Result Stack side by side for presentation (Combined Stack). The Combined Stack was saved as a video file and replayed to review the propagation of SD across hippocampal subfields (Supplementary Video).

Changes in light transmission were quantified using ImageJ. The quantification procedure is illustrated in Supplementary Figure 3. In short, a square box of 50 by 50 pixels was used to sample part of a cell layer of a subfield. The box size was chosen so that it enclosed a representative part of the cell layer with minimal overlap with adjacent layers. Each box was a region of interest (ROI) that was marked and added to the ROI Manager function in ImageJ. These regions included the CA3 cell layer (at or close to the electrophysiological recording site), GCL upper blade, GCL crest, GCL lower blade, and CA1. The Multi Measure function was used to obtain the mean gray value of each ROI (Supplementary Figure 3). Data were visualized in RStudio (v. 1.1.456, http://www.rstudio.com/).

### 2.6 Calculation of SD propagation speed

SD propagation speed was calculated by dividing the length of the SD propagation path (in mm) by the timespan (in min) between two ROIs. An SD propagation path was drawn based on the observations of optical imaging. Because SD propagated from CA3 to CA1 like a wave, and the path followed the curvature of the pyramidal cell layer, the distance between CA3 and CA1 was measured along the curved cell layer and used to calculate the propagation speed. The distance between CA3 and the crest of GCL was measured in a different way because the wave front shown in the intrinsic optical images appeared to first move from CA3 to the upper blade in a linear path and then from upper to lower blade in a path following the curvature of the GCL. Therefore, the distance between CA3 and the crest of GCL was defined as the sum of two segments: (1) a straight line between the origin in CA3 and the tip of upper blade, and (2) a curved line along the GCL from the tip of upper blade to the crest. A timespan between two ROIs were determined by the number of images and the image sampling frequency (every 0.5 or 1 sec).

### 2.7 Quantification

Electrophysiological recordings were quantified using Clampfit software (v. 10.7 or 11.1.0.23, Molecular Devices). Supplementary Figure 1A illustrates how an SD was measured. When an SD was observed, its **onset** was the beginning of the train of bursts just before the DC negative shift. As mentioned above, a burst consisted of at least one population spike superimposed on a positivity. A 5 – 10 sec period before the beginning of the train of bursts was defined as the **baseline**. The **amplitude** of an SD was measured from the baseline to the maximal negative deflection. **Additional synchronized activity** that occurred during the end of the recovery phase, also called afterdischarges, contained negative-going, fast spike-like events in extracellular recordings. This additional synchronized activity was documented in a binary fashion, i.e., slices either had or did not have them. SD **half duration** was the time from the onset of the negative DC shift to the timepoint when the recovery reached ½ of the peak amplitude of the negative DC shift.

Supplementary Figure 1B shows how SLEs were quantified. Similar to an SD, the **onset** of a SLE was the beginning of the train of bursts. There was no large DC negative shift like SD but there could be a small slow DC shift superimposed by bursts during a SLE. The **Amplitude** of this small DC shift during a SLE was defined by the difference between the baseline potential and the most negative deflection. Population spikes were not considered in this measurement. A 5 – 10 sec period before the beginning of the train of bursts was defined as the baseline. The **duration** of a SLE was the onset of the first burst to the time when the last burst ceased.

### 2.8 Statistical analysis

Statistical analyses were performed using Prism (GraphPad) or RStudio (v. 1.1.463, http://www.rstudio.com/). Data were first examined for their normality using the Shapiro-Wilk test. Homogeneity of variance was tested using the F test. An unpaired t test was used to compare means between two groups when data were normally distributed and without significant inhomogeneity of variance. When data were normally distributed with unequal variances, an unpaired t test with Welch’s correction was used. When data violated the assumption of normality, a Mann-Whitney U test was conducted for comparison of two groups. Maximal light change of SD and SLE in CA3 and CA1 was analyzed using a 2-way ANOVA with event type and subregion as between-subject variables. Tukey’s post hoc analysis was followed when the main test reached significance. The incidence of SD and SLE were compared using a Fisher’s Exact test. Differences were considered significant when p < 0.05. Prism, RStudio, Inkscape (v. 0.92, http://www.inkscape.org/), and GNU Image Manipulation Program (GIMP, v. 2.10.18, http://www.gimp.org) were used for preparation of figures.

## 3 Results

### 3.1 An interface-like recording chamber promotes the development of SD and SLE in 0 Mg^2+^/5 K^+^ aCSF

SD in area CA3 occurs in an interface-style chamber when aCSF contains 0 Mg^2+^/5 K^+^ (Mody et al., 1987; Scharfman, 1997). To examine SD in submerged mouse hippocampal slices, we used a classic submerged chamber commonly used in patch clamp recording (Figure 1B). Recordings were made in area CA3 and used extracellular recording methods (Figure 1C) because this was done in prior studies using the interface-style chamber. Many slices (10/16 or 63% of all slices tested, n = 4 mice) did not show any SD or SLE within 60 min of 0 Mg^2+^/5 K^+^ aCSF exposure. However, some slices exhibited SD (3/16) and some showed SLEs (4/16; Table 1).

Because most slices did not exhibit SD or SLEs, and previous reports showed that network activity is facilitated when aCSF flows above and below submerged slices (Hájos and Mody, 2009; Hájos et al., 2009; Morris et al., 2016), we tested an interface-like recording chamber (Figure 1B) using the same slice preparation and recording methods. Figure 1D demonstrates a representative recording in the interface-like chamber. The recording shows the initial period of exposure (specifically from minute 5 to 50 after the start of exposure) of a mouse slice to 0 Mg^2+^/5 K^+^ aCSF. In this example, one SD (arrow) and five SLE (arrowhead) occurred. Typically, we found one SD and many SLEs (discussed further below). Almost all slices developed SD and/or SLEs (44/45 slices, 22 mice). There were significantly more slices that developed SD in the interface-like chamber than in the classic chamber (Fisher’s exact test, p = 0.0009; Table 1).

In prior studies of SD in a submerged chamber, there is often an external stimulus to induce SD such as local application of high potassium or electrical stimulation (Lindquist and Shuttleworth, 2012; Steffensen et al., 2015). To the best of our knowledge, our results are the first to show SD in submerged slices using 0 Mg^2+^/5 K^+^ aCSF.

### 3.2 Characteristics of SD and SLEs in 0 Mg^2+^/5 K^+^ aCSF

#### 3.2.1 SD

Among mouse slices tested in the interface-like chamber 80% of slices developed SD (36/45 slices, 22 mice; Table 1). The majority of slices showed one SD and the most SDs observed were three and it was only found in one slice. The first SD developed after 33.6 ± 1.4 min exposure of 0 Mg^2+^/5 K^+^ aCSF (36 slices, 19 mice). In the slice that developed three SDs, the intervals between onsets of SDs were 12.1 and 13.3 min. The time between the onset of SD and the beginning of the characteristic negative DC shift was 10.7 ± 0.8 sec (36 slices, 19 mice).

The negative DC shift consisted of one or two peaks. Seventy-seven percent of slices that developed SD showed two peaks (27/35 slices, 18 mice; 1 slice was not included due to a technical problem that occurred during the recording). The time from the beginning of the negative DC shift to the first peak was 2.2 ± 0.2 sec and to the second peak was 22.6 ± 1.9 sec (27 slices, 18 mice). The peak amplitude of the first and the second peaks were −3.1 ± 0.4 and − 3.5 ± 0.4 mV, respectively (27 slices, 18 mice). The amplitude difference between two peaks was mostly within one standard deviation with only three exceptions, where two slices had a larger second peak and one slice had a larger first peak. For slices that developed only one SD, the timespan from the beginning to the peak of negative DC shift was 22.6 ± 12.7 sec (8/35 slices, 7 mice).

The half duration of SD was 53.0 ± 4.7 sec (35 slices, 19 mice). During the recovery phase, 29/36 slices showed additional synchronized activity. The synchronized activity lasted for 102.9 ± 6.1 sec (29 slices, 17 mice). No sex difference was observed in the measurements of incidence of SD, number of SD, SD onset, and maximal SD amplitude (Supplementary Figure 2).

In a subset of experiments, simultaneous whole-cell recording was performed to examine the activity of individual CA3 pyramidal cells (PCs) during SD (5 cells, 5 mice; Figure 2A). Recorded cells were filled with biocytin, and the cell type was confirmed later to have the morphological characteristics of CA3 PCs, such as a prominent apical dendritic tree, basal dendrites, and spiny dendrites (Lorente De Nó, 1934; Figure 2B). The electrophysiological characteristics also were consistent with a PC, such as an AP with a time course similar to a “regular spiking” neuron and an absence of the characteristic large afterhyperpolarizations of GABAergic neurons (Scharfman, 1993b, 1993a, 1995).

**Figure 2.**
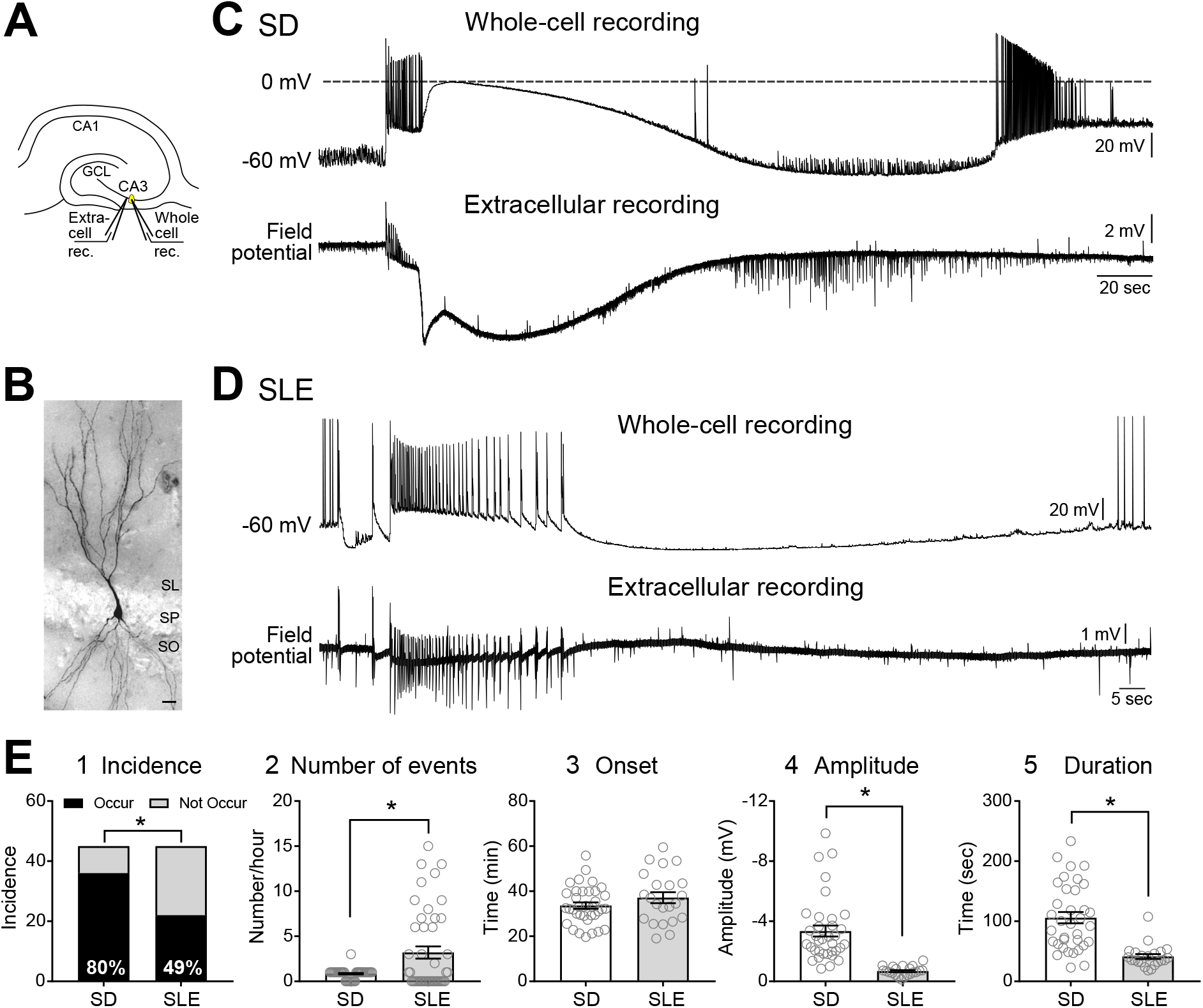
SD and SLEs are two main types of events with distinct characters in 0 Mg^2+^/5 K^+^ aCSF slice model. **(A)** Schematic illustrates the recording sites for simultaneous field potential and whole-cell recordings in CA3. **(B)** A representative image of a recorded and biocytin-filled CA3 pyramidal cell. SL, stratum lucidum; SP, stratum pyramidale; SO, stratum oriens. Scale bar, 20 µm. **(C)** Example of an SD. The intracellular and extracellular characteristics of SD are shown. **(D)** Example a SLE. The intracellular recording shows characteristics of a SLE such as a sudden depolarization and extensive firing. **(E)** Comparisons between SD and SLEs. (**E1)** In a 60-min recording period, SD showed higher incidence than SLEs. Two-sided Fisher’s exact test (p = 0.02). **(E2)** In a 60-min recording period, more SLEs occurred than SD. Wilcoxon test (p = 0.01). **(E3)** The onset of the first SD and SLE was not significantly different. Unpaired t test (p = 0.18). **(E4)** SD triggered a larger negative deflection than SLEs. Mann-Whitney test (p < 0.0001). **(E5)** SD had a longer duration than SLEs. Mann-Whitney test (p < 0.0001). No sex difference was observed (Supplementary Figure 2).

In the 5 recorded PCs, a sudden, large depolarization that occurred at the start of SD was 21.5 ± 2.4 mV. After the cessation of cell firing, a second larger depolarization of 39.5 ± 3.7 mV followed. Therefore, the peak of the depolarizations was close to 0 mV. As expected for SD, the depolarization phase corresponded to the negative DC shift observed in the extracellular recording (Figure 2C). Cells repolarized and then hyperpolarized for 7.4 ± 3.2 mV before returning to their resting membrane potentials. Additional synchronized activity observed in extracellular recordings corresponded to large amplitude depolarizations and/or AP firing (Figure 2C).

#### 3.2.2 SLEs

In 45 tested mouse slices, 49% of slices developed SLEs within the 60 min of exposure to 0 Mg^2+^/5 K^+^ aCSF (22/45 slices, 22 mice; Table 1). Among 22 slices that showed SLEs, 8 of them developed only SLEs and the other 14 slices developed both SD and SLEs. More SLEs developed in slices without an SD (without SD, 8 slices, 10 ± 1 SLEs; with SD, 14 slices, 5 ± 1 SLEs; unpaired t test, t (20) = 2.743, p = 0.01), suggesting a negative interaction between SD and SLEs.

The average duration of SLEs was 41.4 ± 4.0 sec (22 slices, 14 mice). Although the duration of SLEs was indifferent with or without an SD (without SD, 8 slices, 42.8 ± 2.8 sec; with SD, 14 slices, 40.7 ± 6.2 sec; unpaired t test with Welch’s correction, t (17.42) = 0.3069, p = 0.76), SLEs with SD showed greater variation than SLEs without SD (F test, F = 8.8, p = 0.0079). This result supports the idea mentioned above that SDs interfere with SLEs.

During SLEs, an extremely small negativity developed. The average negativity was −0.7 ± 0.1 mV (21 slices, 14 mice). The average negativity of SLEs did not appear to depend on appearance of SD (without SD, 8 slices, −0.7 ± 0.1 mV; with SD, 13 slices, −0.7 ± 0.1 mV; unpaired t test, t (19) = 0.2455, p = 0.81).

For 7 cells in 5 mice, intracellular recordings were made simultaneous to extracellular recordings and showed the components of SLEs (Figure 2D). The trains of bursts during SLEs were qualitatively similar to the initial train of bursts at the onset of SD (Figure 2C). However, the duration of the train of bursts of an SLE was longer than the train of bursts at SD onset (SLEs, 40.9 ± 4.2 sec, 21 slices, 13 mice; train of bursts at SD onset, 9.3 ± 0.8 sec, 33 slices, 18 mice; Mann-Whitney test, U = 3, p < 0.001). The similarity in the trains of bursts of SLEs and SDs suggest that an SLE was a failure of bursts to trigger SD.

#### 3.2.3 Comparisons between SD and SLEs

##### 3.2.3.1 Incidence

During the first 60 min of exposure to 0 Mg^2+^/5 K^+^ aCSF, 31% of mouse slices developed both SD and SLE (14/45 slices, 22 mice) while 49% of slices developed only SD (22/45 slices), and 18% of slices developed only SLEs (8/45 slices). The differences were significant, with more slices exhibiting SD than SLE (SD, 36/45 slices; SLE, 22/45 slices, Fisher’s exact test, p = 0.0039, Figure 2E1). Each slice developed significantly fewer SDs than SLEs (SD, 0.8 ± 0.1; SLE, 3.2 ± 0.7; 45 slices, Wilcoxon matched pairs signed rank test, p = 0.049; Figure 2E2).

##### 3.2.3.2 Onset of SD and SLE

The first SD occurred 33.6 ± 1.4 min after the slice was transferred into the recording chamber with 0 Mg^2+^/5 K^+^ aCSF (36 slices, 22 mice), which was not different from the first SLE (37.1 ± 2.4 min, 22 slices, 21 mice; unpaired t test, t (56) = 1.348, p = 0.18; Figure 2E3). Among 14 slices that showed both SD and SLEs, SD was more likely to develop before a SLE (SD as the first event, 11/14 slices, SLE as the first event, 3/14 slices; Fisher’s exact test, p = 0.0027). Thus, the onset of SD and SLE was similar, but an SD was more likely to be the first event in our 0 Mg^2+^/5K^+^ aCSF preparation.

##### 3.2.3.3 Amplitude and half-duration

When the DC shift was measured (for Methods, see Supplementary Figure 1), SDs showed an average of a 4.7 times larger negative shift than SLEs (SD, −3.3 ± 0.4 mV, 36 slices, 19 mice; SLEs, −0.7 ± 0.1 mV, 22 slices, 14 mice; Mann-Whitney test, U = 12, p < 0.0001; Figure 2E4). When an SD’s duration was estimated by doubling SD’s half-duration, SD had duration more than twice the duration of SLEs (SD, 106 ± 9 sec, 36 slices, 19 mice; SLE, 41 ± 4 sec, 22 slices, 14 mice; Mann-Whitney test, U = 12, p < 0.0001, Figure 2E5). No sex difference was observed in these measures (Supplementary Figure 2).

### 3.3 SD propagation

#### 3.3.1 SD spread to CA1 vs. dentate gyrus

Propagation across gray matter is an important feature of SD. When 0 Mg^2+^/5 K^+^ aCSF was used in the past, it was suggested that SD usually begins in CA3 and then spreads to CA1 in hippocampus (Mody et al., 1987). Other studies using rabbit hippocampal slices and focal K^+^ to elicit SD came to a similar conclusion (Haglund and Schwartzkroin, 1984). A spread to the dentate gyrus through upper blade has been reported when SDs are induced in CA1 using a local high-K model (Buchheim et al., 2002). However, spread from CA3 to the dentate gyrus has not been studied before using 0 Mg^2+^/5 K^+^ aCSF. Therefore, we first confirmed that the SDs we observed did spread and then asked how they spread from CA3.

##### 3.3.1.1 Extracellular recordings

We began by electrophysiological recording in two locations using extracellular methods. First, we examined the propagation of SD from CA3 to CA1 in 0 Mg^2+^/5 K^+^ aCSF. Extracellular recording electrodes were placed in the CA3b and CA1b regions (Figure 3A1). In seven slices, five developed an SD in CA3. In two of these slices, CA1 showed an SD. Consistent with past reports, the onset of the negative DC shift in CA1 was delayed from CA3 by an average of 8 sec (range 5 – 11 sec, Figure 3A2) whereas the initial train of bursts occurred at similar time (< 10 msec). These data are interesting because they suggest neuronal activity can occur with short delays but the drastic cell depolarization, occurring during the negative DC shift, is first in CA3 and very slowly propagates to CA1.

**Figure 3.**
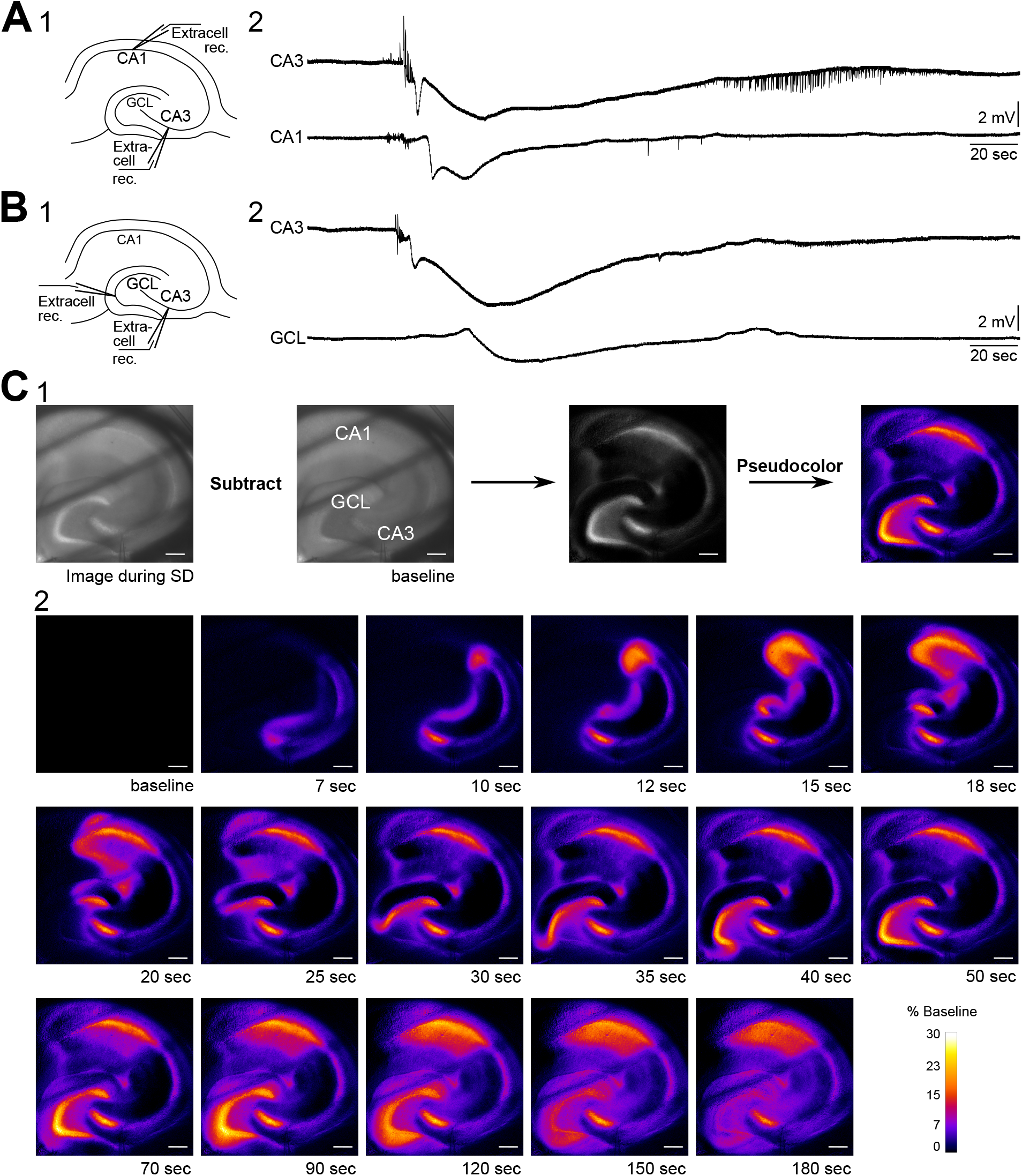
Propagation of SD in mouse hippocampus. **(A1)** Schematic of dual field recordings in the CA3 and CA1. **(A2)** An SD propagated from CA3 to CA1. The electrical recordings show a significant delay between the large negativity, corresponding to the delay in the optical image in C. **(B1)** Schematic of dual field recordings in the CA1 and GCL. **(B2)** An SD propagated from CA3 to GCL with a time delay. **(C1)** An SD was accompanied by a changed in the intrinsic optical signal. The difference in light transmission was calculated by subtracting a baseline image from the image acquired during SD. **(C2)** Consecutive images illustrate the propagation of an SD. Diagonal dark lines were from the mesh underneath the brain slice. GCL, granule cell layer. Scale bar, 200 µm.

Next, we examined the propagation of SD from CA3 to the dentate gyrus. Extracellular recording electrodes were placed in the CA3 PCL and GCL at the crest of the dentate gyrus, which was chosen because it is the furthest point from CA3 in the GCL (Figure 3B1). In two slices with these recording locations, both slices showed SD in the GCL with a long delay of 26.7 sec (range 24.9 – 28.5 sec, 2 slices from 2 different mice). Our data suggest that SD that develops in CA3 during superfusion of 0 Mg^2+^/5 K^+^ aCSF can propagate to both CA1 and dentate gyrus and with different delays, and the delay from CA3 to the dentate gyrus appears to be longer than the delay from CA3 to CA1.

##### 3.3.1.2 Intrinsic optical imaging

SD-induced neuronal swelling changes the light transmission. Specifically, the light transmission increases when an upright brightfield microscope is used (Figure 3C1, Buchheim et al., 2002; Mané and Müller, 2012). We used this characteristic of SD to examine the path of the SD wavefront across hippocampal subregions in our slice preparation.

To capture changes in light transmission that reflected SD propagation, images of SDs were taken and subtracted from the baseline image (see Methods and Figure 3C1). Figure 3C2 shows images selected in temporal order throughout an SD. In 9/11 imaged slices the wave fronts of the SDs all emerged from CA3 (Figure 4). SDs of the exceptional two slices were not fully imaged so the emergence of their wavefronts was unclear. Within the 9 slices, 44% of SDs propagated into both CA1 and the whole dentate gyrus (4/9 slices). Other SDs propagated either only into the whole dentate gyrus (2/9 slices), or only into hilus (2/9 slices), or only into CA1 (1/9 slices). The direction of SD propagation was either toward CA1 or toward dentate gyrus or both. No association between propagation pattern and dorsoventral axis of hippocampus was observed.

**Figure 4.**
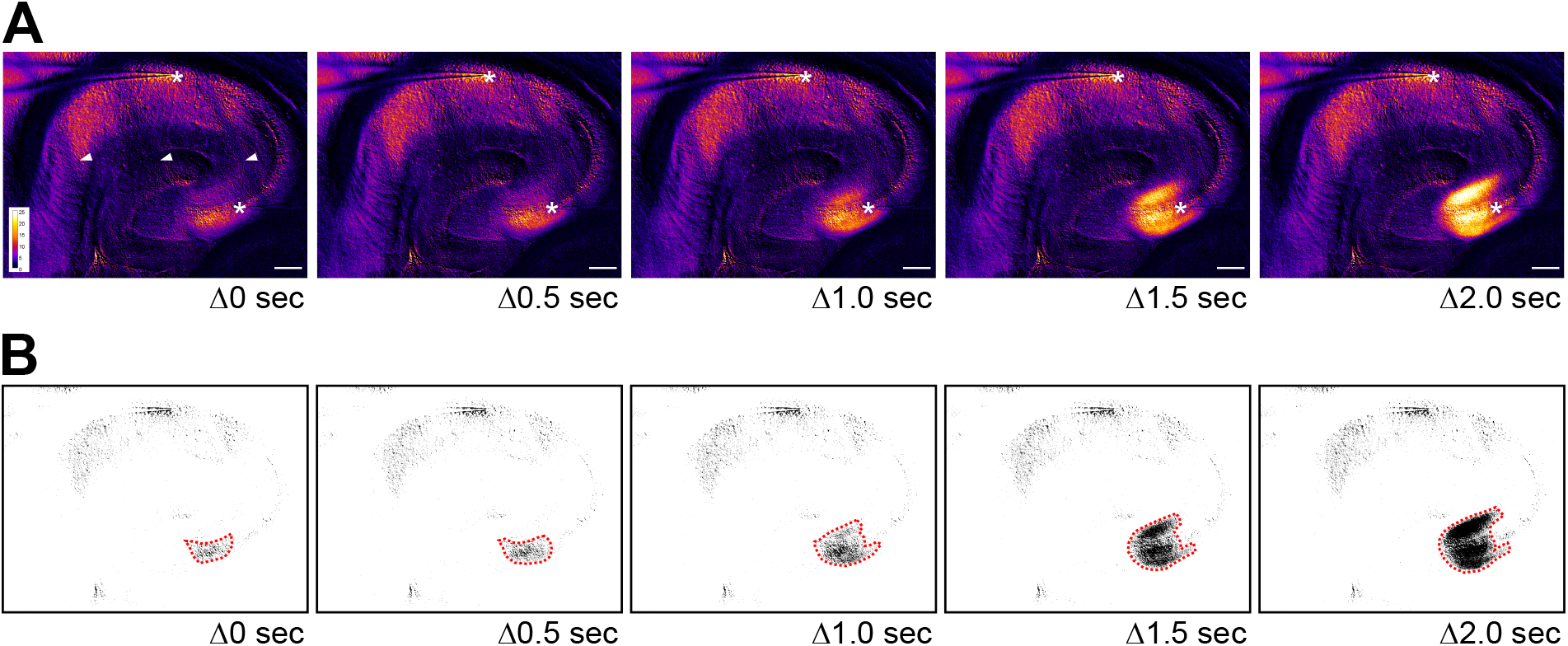
The initial wave of spontaneous SD often begins in CA3 in 0 Mg^2+^/5 K^+^ aCSF. **(A)** Time lapse images show the initiation site of an SD in CA3. The images during SD were first subtracted from a baseline image and then pseudocolored. Warmer color indicates higher light transmission and cooler color for lower light transmission. The increase of light transmission began in both SP and SL of CA3 before a wave front became prominent. Stars: recording sites in CA3 and CA1. Arrowheads: dark areas reflecting the mesh underneath the slice. Scale bar, 200 µm. **(B)** Same images from panel A were thresholded so that area with the greatest change in light transmission in CA3 could be distinguished (dashed red lines).

To quantify the propagation speed of SDs, we measured the length of the SD traveling path among our defined regions of interests (see Section 2.6) and the time that elapsed from the start of the path to its end. The propagation speed from CA3 to CA1 was 4.4 ± 1.6 mm/min (range 2.6 – 8.6 mm/min, 5 slices, 3 mice). The propagation speed from CA3 to the crest of the GCL in the dentate gyrus was 2.7 ± 0.5 mm/min (range 1.8 – 4.6 mm/min, 9 slices, 4 mice). Propagation speed from CA3 to CA1 was not statistically different from the speed from CA3 to GCL (Unpaired t test with Welch’s correction, t (4.706) = 2.284, p = 0.07).

Supplementary Video shows the emergence, propagation, and recovery of an SD using original brightfield and pseudocolored images. Taken together, these results show that most SD wave fronts emerge from CA3 in 0 Mg^2+^/5 K^+^ aCSF. SD then spreads both in the direction of the trisynaptic circuit, i.e., to CA1, and in the opposite direction of the trisynaptic circuit, i.e., to the dentate gyrus. The speeds are similar and consistent with past studies (Obeidat and Andrew, 1998; Buchheim et al., 2002).

#### 3.3.2 SD spread from upper to lower blades of the dentate gyrus

When SDs in CA3 propagated toward the dentate gyrus, 25% of SDs (3/12 slices, 6 mice) reached the hilus and/or upper blade of GCL but then stopped spreading to the other parts of the dentate gyrus. The rest of the SDs (9/12 slices, 6 mice) traveled to the upper blade and then spread to the crest and finally the lower blade (Figure 5A).

**Figure 5.**
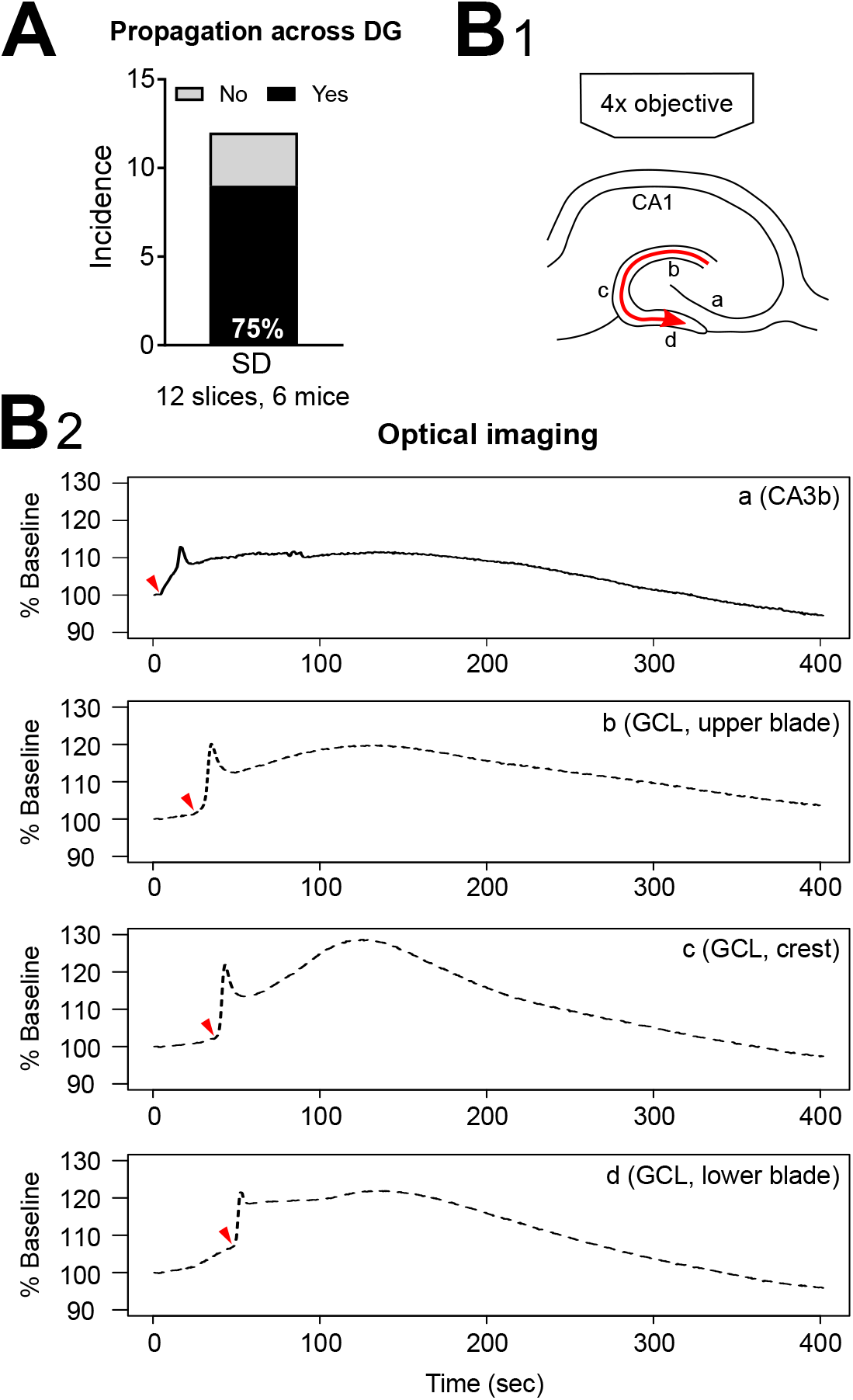
Propagation of SD within dentate gyrus. **(A)** SD propagated from upper to lower blades occurred in the majority (77.8%) of optically recorded slices. **(B1)** A schematic of SD propagation. The red arrow indicates the observed direction of SD propagation. **(B2)** Representative traces of light changes in CA3b (a), GCL, upper blade (b), GCL, crest (c), and GCL, lower blade (d). Red arrowheads indicate the time when light transmission began the most rapid phase of the onset, showing the propagation of SD schematized in B1.

SDs we observed entered dentate gyrus in two separate waves: into the hilus and into the tip of upper blade (Figure 3C2). Although the lateral tip of lower blade of GCL was also approached by the wave front (Figure 4), it was not clear that this was a separate path from entry into the hilus, which appeared to spread into the entire area between both blades at once. The wave front stopped at the boundary of hilus with the GCL. The wave front that entered the upper blade moved through the GCL and molecular layer together and followed the curvature of the GCL so that it traveled from upper blade to crest to lower blade in a slow continuous wave (Figure 5B1).

Quantification of light transmission supported the propagation pattern of SD wave front. The onset of sharp rise of light transmission happened in sequence among CA3b (Figure 5B2a), GCL upper blade (Figure 5B2b), GCL crest (Figure 5B2c), and GCL lower blade (Figure 5B2d). After the first peak in light transmission there was a short drop of light transmission followed by an increase of light transmission in the GCL and molecular layers (Figure 5B2, Supplementary Video). This secondary increase in light transmission was also observed in CA3 and CA1, and the underlying cellular correlates remain unclear.

To quantify the speed of propagation within the dentate gyrus, we measured the length of SD propagation by following SD’s traveling path, and the time spent from upper blade, crest to lower blade (see Section 2.6). The speed of propagation was 2.8 ± 0.5 mm/min (range 1.4 – 4.0 mm/min, 7 slices, 4 mice).

### 3.4 SLEs showed increased but weak intrinsic optical signals

Next, we quantified changes in light transmission of SLEs. Figure 6 compares the change in light transmission of SDs to SLEs. In contrast to SD (Figure 6A2), the change in light transmission during the SLEs was minimal (Figure 6B2). However, although changes in light transmission in SLEs were small, they were significantly greater than baseline (One sample t test, CA3, t (6) = 4.75, p = 0.0032; CA1, t (6) = 3.96, p = 0.0075).

**Figure 6.**
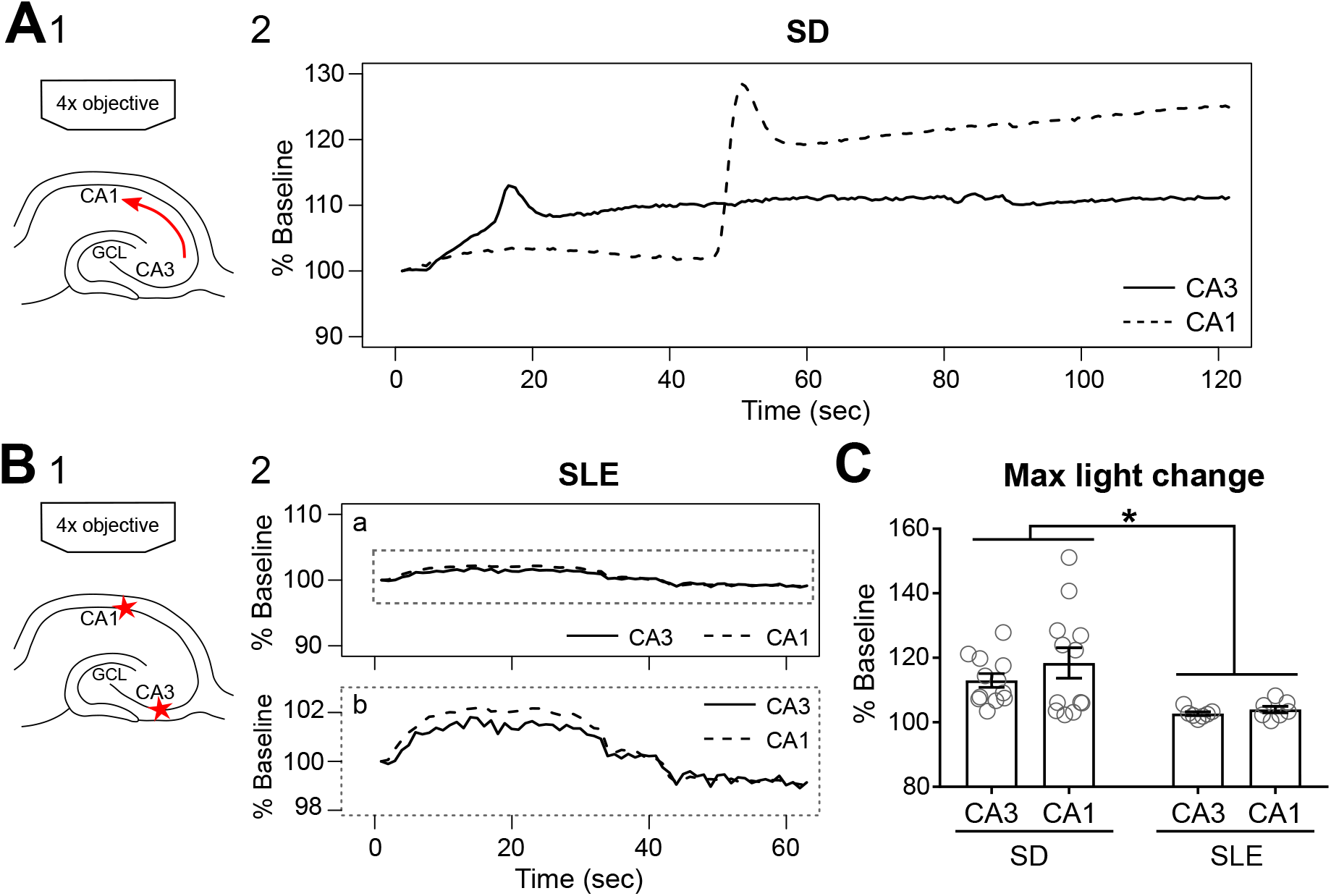
SDs, but not SLEs, show prominent changes in light transmission. **(A1)** Schematic of the path of light transmission in A2. Arrow indicates the propagation of the observed SD. **(A2)** Change of light transmission initiated in CA3 (solid line) and then followed by CA1 (dashed line) with a delay. **(B1)** Schematic of the region of interests (stars) in B2. **(B2)** A SLE showed subtle changes in light transmission (expanded in b) without a long delay between CA3 (solid line) and CA1 (dashed line). **(C)** SDs had significant higher change in light transmission than SLEs (Two-way ANOVA, SD vs. SLEs, F (1,34) = 12.69, p = 0.0011; CA3 vs. CA1, F (1,34) = 0.34, p = 0.3376; interaction, F (1, 34) = 0.3585, p = 0.5533).

It was not surprising that the percent change of maximal light transmission of SD was greater than SLEs (Two-way ANOVA, SD vs. SLEs, F (1,34) = 12.69, p = 0.0011; CA3 vs. CA1, F (1,34) = 0.34, p = 0.3376; interaction, F (1, 34) = 0.3585, p = 0.5533; Figure 6C). This was true for CA1 (SD, 118.4 ± 4.7%, 12 SDs in 12 slices, 6 mice vs. SLEs, 104.0 ± 1.0%; 7 SLEs in 7 slices, 3 mice; p = 0.03, Tukey’s multiple comparison test) but not for CA3 (SD, 113.0 ± 2.1%, 12 SDs in 12 slices, 6 mice vs. SLEs, 102.7 ± 0.6%; 7 SLEs in 7 slices, 3 mice; p = 0.18, Tukey’s multiple comparison test). These data suggest that electrical recordings are quantitatively more sensitive than light transmissions.

In summary, SLEs showed a very small but detectable change in light transmission. The data are consistent with studies showing an increase in light transmission associated with an increase in neuronal activity (MacVicar and Hochman, 1991; D’Arcangelo et al., 2001).

## 4 Discussion

### 4.1 Summary

In the present study we first addressed methods to use mouse slices to study SD and SLEs during exposure to 0 Mg^2+^/5 K^+^ aCSF. We recorded these events using extracellular recording, whole-cell recording, and intrinsic optical imaging, which was advantageous both temporarily and spatially. We found that the chamber mattered: SD was more likely to develop in submerged mouse slices in an interface-like than in a classic recording chamber. Species also mattered: in the submerged condition, mouse hippocampal slices developed SD and rat slices did not show SD. However, both species showed SLEs. Notably, recording and imaging both SD and SLEs provided a chance to examine them in ways that have rarely been done in the past.

Intrinsic optical imaging allowed us to examine the emergence of the SD wave front and SD propagation without investing in additional dyes, transgenic mice, or equipment. Within hippocampus, we found that the majority of SDs initiated in CA3. Because of our imaging field only covered the hippocampus with limited cortical areas, we cannot exclude that SDs initiated in the small part of the cortex attached to the hippocampus (Martens-Mantai et al., 2014). Interestingly, when we recorded extracellular electrical activity in CA3 and CA1 simultaneously, the SDs began with a train of bursts with less than 10 msec delay between CA3 and CA1. However, the negative DC shift in CA3 immediately followed the train whereas in CA1 it was clearly delayed. The concurrent trains of bursts may have propagated synaptically from CA3 to CA1 but the drastic cell depolarization that underlies the DC shift relies on other mechanisms. Alternatively, the entorhinal cortex innervates both CA3 and CA1 pyramidal cells (Witter et al., 2017) and may trigger trains of bursts.

In hippocampus, SD propagated along the direction of classic trisynaptic circuit into CA1 and in the opposite direction into dentate gyrus. Among SDs that propagated into dentate gyrus, they traveled in a way that is not consistent with the idea that SD spreads in all directions by a simple process. Instead, SDs propagated in two separate waves—one moved into hilus and the other moved along molecular and granule cell layers. The wave front that propagated into hilus obeyed the boundary between granule cell layer and hilus, and it dissipated in hilus. The wave front that moved into molecular and granule cell layers propagated from upper blade to crest, and then lower blade. The wave front encompassed the entire molecular layer without involving adjacent layers in CA1 or the hilus. Although other researchers have similar observations using local high K^+^ injections to initiate SD in CA1 (Obeidat and Andrew, 1998; Buchheim et al., 2002), our report is the first one to show the unique SD propagation pattern in dentate gyrus using 0 Mg^2+^/5 K^+^ aCSF. The consistent direction of SD propagation across different SD induction models suggests that a common structural factor, e.g., the components of the neuropil like the extracellular matrix and vasculature which is not necessarily homogeneous. In particular, the GCL/hilar border has a very different milieu, which makes it subject to injury (Soltesz et al., 1995).

In addition to SD, SLEs developed in 0 Mg^2+^/5 K^+^ aCSF. SD and SLE had distinct characteristics: (1) SDs’ negative DC shift were more than 5x that of SLEs’; (2) SDs were approximately 6x longer than SLEs; and (3) SDs had approximately 7x larger changes in light transmission than SLEs. However, the trains of bursts underlying SLEs were quite similar to those before SD. These results suggest that once a SLE is initiated in CA3, it could complete as a SLE or trigger an SD. Possible factors that could switch the progression of a SLE into an SD may be related to the degree the slice preparation preserves the recurrent collaterals of CA3 PCs, which would allow greater synchrony and in turn, greater local accumulation of ions. The mossy fiber pathway may also be a factor, because it can vary in the degree it is maintained in a slice also; if well maintained its normally high concentration of glutamate can be released onto CA3 PCs to a greater extent, facilitating SD. Although our data do not suggest major variations in these factors, slices can vary from one to the next and from one animal to another.

Taken together, our optimized model with 0 Mg^2+^/5 K^+^ aCSF led to a slow development of both SD and SLEs, which provides an opportunity to study these events without the typical trigger where the stimulus is immediately followed by SD. Additionally, our preparation also provided great temporal and spatial resolution by combining electrophysiological recording and optical imaging. We therefore think that this experimental model has many uses to advance our understanding of mechanisms of SD and interactions between SD and SLEs. Also, one could use the approach for validating preclinical drug candidates by taking advantage of a controlled ex-vivo environment and relevance to disease mechanisms of migraine, traumatic brain injuries, stroke, etc.

### 4.2 The importance of the recording chamber

Submerged slices seem to be a suboptimal condition for SD and even SLE development when compared to slices situated at an interface of oxygenated solution and air. Thus, slices maintained at an interface, from rats showed both SD and SLEs in 0 Mg^2+^/3 – 5 K^+^ aCSF (Mody et al., 1987; Gloveli et al., 1995; Kovács et al., 1999), but only individual bursts and SLEs were reported in submerged rat hippocampal slices in 0 Mg^2+^/2 – 5 K^+^ aCSF (Lewis et al., 1989, 1990; Neuman et al., 1989; Churn et al., 1991; Kojima et al., 1991; Billard et al., 1997; Wahab et al., 2009). When rat slices in a classic submerged chamber were compared with rat slices in an interface chamber, using the same experimental conditions, fewer submerged slices show SLEs than slices in the interface chamber (Schuchmann et al., 2002). Moreover, anoxia induced SD only in rat slices recorded in an interface-style chamber but not in a classic submerged-style chamber (Croning and Haddad, 1998). Our data are consistent with these findings.

Both SD and SLEs are types of synchronized activity. An in vitro environment that supports synchronized activity would aid the development of SD and SLEs, and oxygen supply could be a critical player. The oxygen in interface-style chambers is supplied from carbogen-saturated humidified air and aCSF, while carbogen-saturated aCSF is the sole oxygen resource in submerged-style chambers. Since only a limited amount of oxygen can dissolve in aCSF, the direct contact with humidified air in an interface-style chamber has been considered as a more efficient way to supply oxygen than slices that obtain all oxygen from aCSF in a submerged-style chamber (Aitken et al., 1995). In interface-style chambers, rat brain slices showed in vivo-like rhythmic oscillations generated by synchronized cell activity (von Krosigk et al., 1993; Sanchez-Vives and McCormick, 2000; Case and Broberger, 2013). In submerged-style chambers, however, similar observations of synchronized activity seem possible only when oxygen supply is increased by superfusing aCSF above and below slices and using a fast flow rate (Hájos and Mody, 2009; Hájos et al., 2009). We found that the likelihood of developing SD in mouse slices significantly increased in an interface-like chamber, where 80% of mouse slices developed SD in an interface-like recording chamber when compared to 19% of slices in a classic recording chamber. Our results agree with previous findings and support the hypothesis that oxygen supply is important in SD and SLEs.

### 4.3 Species differences in the development of SD in 0 Mg^2+^/5 K^+^ aCSF

When mouse and rat slices were compared in the same preparation (interface-like chamber, fast flow, and 0 Mg^2+^/5 K^+^ aCSF), we found that SLEs were observed in slices from both species while SDs were exclusively in mouse slices, at least for SDs and SLEs following 0 Mg^2+^/5 K^+^ aCSF exposure. The discrepancy may be explained by the preservation of circuit components. We prepared both mouse and rat hippocampal slices using the same thickness (350 µm). Using the same thickness, it has been proposed that mouse slices are likely to preserve more connections and circuit properties than rat slices (Heinemann et al., 2006). It is also possible that the rat slices needed more oxygen because they were larger, despite the same thickness as the mouse slices. Another possibility to explain the species difference could be related to age. The ranges of ages overlapped but the youngest mice were younger than the youngest rats. On the other hand, we tested very young rats and mice (postnatal day 21 – 22) and we obtained similar results—rat slices did not exhibit SD, but mouse slices did. Therefore, age did not appear to be a factor in species differences.

### 4.4 SD vs. SLEs in 0 Mg^2+^/5 K^+^ aCSF

In slices that developed both types of events, 73% of slices developed SD first. Most slices exhibited only one SD (32 of 33 slices; the exceptional slice had 3 SDs) in the first hour of 0 Mg^2+^/5 K^+^ aCSF incubation. On the other hand, a slice could show multiple (1 – 16) SLEs with or without developing SD. These results are consistent with the suppression of neural activity after SD. For example, in rat slices in an interface-style chamber, SDs occur spontaneously after 0 Mg^2+^/5 K^+^ aCSF exposure but only every 15 minutes. For minutes after an SD, responses to afferent stimulation in CA3 were weak (Scharfman, unpublished).

Both SD and SLEs developed in CA3 of mouse hippocampal slices in 0 Mg^2+^/5 K^+^ aCSF. When we examined the onset of SD, we found trains of bursts that were very similar to SLEs, but shorter. Why these bursts are followed by SD sometimes, but otherwise they do not appear to do so, is not clear. It could be the bursts are triggered by different mechanisms, one due to entorhinal cortical input and one intrinsic to CA3. These different mechanisms may recruit feedforward or feedback inhibition more for SLEs and less for SD. Another possibility is that SLEs sometimes trigger SD and sometimes do not. This hypothesis is consistent with the idea that SD can be a natural anti-seizure mechanism where a seizure triggers SD and then SD suppresses activity for some time afterwards (Tamim et al., 2021).

### 4.5 Propagation of SD

We found that SD typically began in CA3. We used two methods to identify this: electrical recording from more than one location and intrinsic optical imaging. Since our imaging field was limited, our method did not allow us to determine if SD began in an extrahippocampal area but use of a lower power objective could do so in the future. An increase of light transmittance propagated toward both CA1 and dentate gyrus with up to 30% transmittance change locally. Interestingly, SLEs only showed subtle changes in the intrinsic optical signals (2%) but it was possible to visualize the small changes. It is possible that more detail might be clear with other wavelengths, such as 460 – 560 nm used previously (Mané and Müller, 2012). However, our recording at 775 nm was certainly sufficient.

Our analysis of SD propagation to the dentate gyrus is one of the few studies that have examined this issue. We found that SD propagated toward dentate gyrus in a unique pattern. Instead of moving like a unidirectional wave, which approached the upper and lower blade simultaneously before the wave front reached crest, SD circled around the dentate gyrus through upper blade, crest, and then lower blade. Our data are consistent with a previous finding when focal K^+^ application to CA1 was used (Buchheim et al., 2002).

### 4.6 Advantages and disadvantages of our methods

There are several advantages to the methods that we used. Our slice model provides a method to expand the repertoire of a patch clamp rig to study spatiotemporal dynamics of both SD and SLEs without costly investments. While we found limitations with the classic submerged-style chamber, a slight modification to use a design more like an interface-style chamber circumvented the problem. This interface-like design can be fabricated or, as was the case in our study, purchased. The cost of a chamber was one of the few investments our methods required. Using a fast flow rate seemed important and is highly feasible.

Although rat slices could be used with the interface-like chamber, SD was not common. This might be circumvented by a mini-slice (if the problem is the large size of the slice, mentioned above), but then all subfields could not be imaged. Therefore, rat studies of SD are feasible with our methods but will require more animals to sample SD enough for statistical comparisons.

One advantage of our methods was that SD occurred faster than in prior studies using a classic interface-style chamber. SD only spontaneously developed after 2 hours of 0 Mg^2+^/5 K^+^ aCSF exposure in a previous study (Mody et al., 1987). In comparison, we observed spontaneously developed SD within the first hour of 0 Mg^2+^/5 K^+^ aCSF incubation.

The methods can potentially be used with ways to induce SD other than 0 Mg^2+^/5 K^+^ aCSF, such as local high K application, low calcium, ouabain, etc. It also can be used with transgenic mice, optogenetics, chemogenetics, and other methods currently used in research. Pharmacology is more readily used with the submerged chamber designs due to a smaller volume in the chamber compared to classic interface chambers. Therefore, drug studies and preclinical studies are facilitated by our methods.

The ability to induce both SD and SLEs is also useful for future studies of neuronal excitability and preclinical studies for epilepsy. For example, when a drug will be inhibitory to SD but facilitate SLEs is important to identify. The opposite, a drug that inhibits SLEs but leads to more SD is important.

Although we found no sex differences in the development of SD in mouse hippocampal slices. This may simply be due to the fact that the mice had not yet reached sexual maturity (Bell, 2018). Extending our findings into adulthood will be valuable in the future because of sex differences and hormonal regulation of in SD (Scharfman et al., 2003; Harte-Hargrove et al., 2013; Skucas et al., 2013; Scharfman and MacLusky, 2014). This topic has important translational implications because of sex differences in migraine and epilepsy (Scharfman and MacLusky, 2008, 2014; Vetvik and MacGregor, 2017).

There are also some disadvantages to the methods that we used. Although we were able to examine SD without a focal stimulus, the unpredictability of these SDs made it hard to trigger the start of video at the baseline before SD occurred. One method to circumvent the issue is to record throughout exposure to 0 Mg^2+/^5 K^+^ aCSF, and as digital acquisition and storage improve such prolonged recording is likely to be easier.

## Conclusion

SD and SLEs are both products of hypersynchronized activity of brain cells. We optimized a classic 0 Mg^2+^/5 K^+^ aCSF model for submerged hippocampal slices so both SD and SLEs are spontaneously developed within the first hour of 0 Mg^2+^/5 K^+^ aCSF exposure. We also extended our ability to record intrinsic optical signals without additional costs. With the relevance of SD and seizure activity to many pathological conditions and neurological disorders, this optimized model provides an opportunity to study SD with temporal and spatial advantages. It could also address how SD and SLEs interact with each other in a controlled environment. We suggest that this model could be useful in testing mechanisms and validating treatment targets in pre-clinical studies.

## Author contributions

Y-L.L. and H.E.S. conceived and designed the study. Y-L.L. performed the experiments and analyzed the data. Y-L.L. and H.E.S. wrote the manuscript.

## Acknowledgements

This study was supported by NIH grants AG-055328, MH-109305, NS-037562, NS-081203, and Pyramid Biosciences. We thank Karim Elayouby for the technical support in immunolabeling experiments. We are grateful to (by alphabetical order) Elissavet Chartampila, Kasey Gerencer, Swati Jain, John LaFrancois, and Drs. David Alcantara-Gonzalez, Hannah Bernstein, Justin Botterill, Chiara Criscuolo, Áine Duffy, and Christos Lisgaras for their experimental supports and valuable discussions and feedback on the manuscript.

---

**Supplementary Figure 1.**
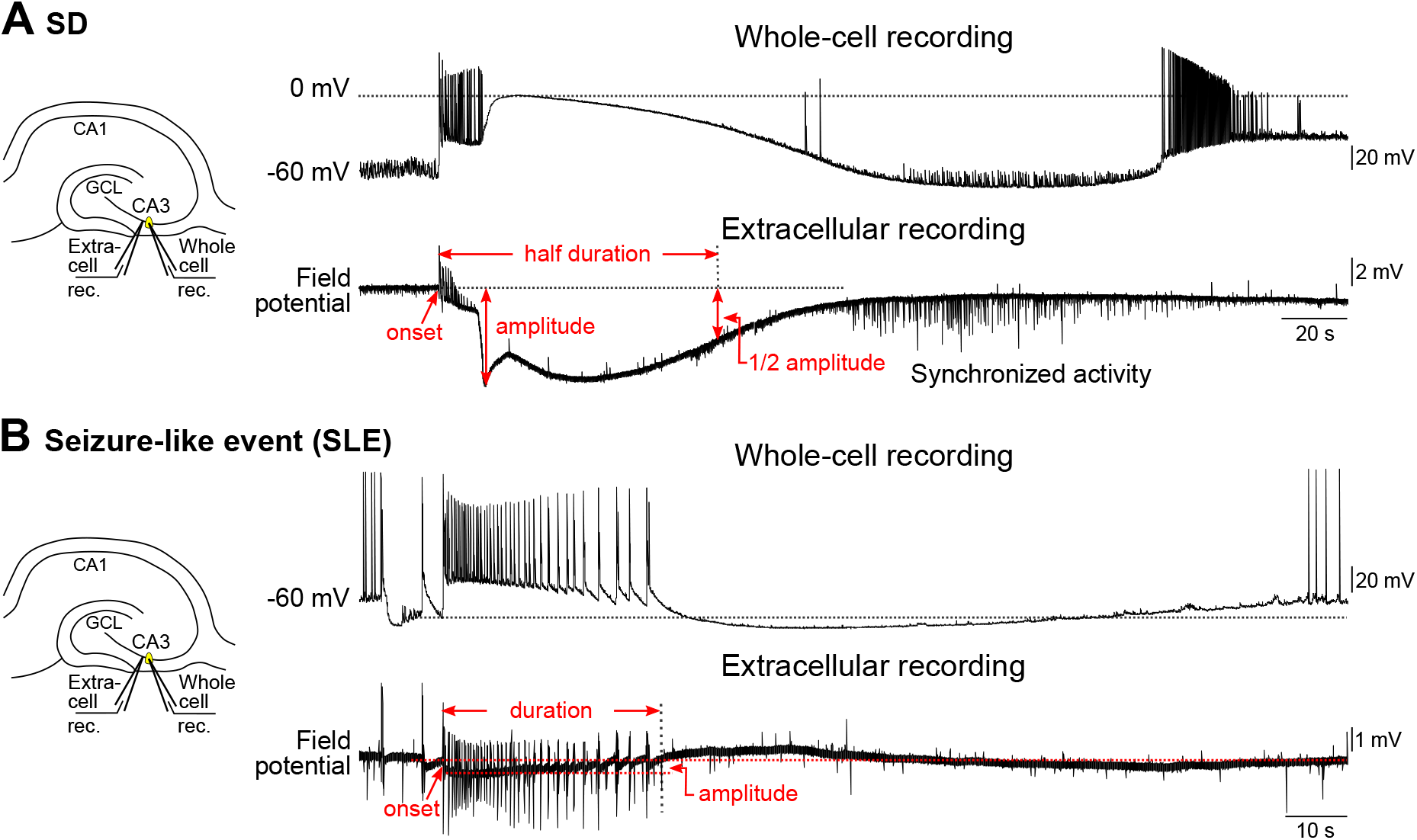
Schematic illustrates the quantification for SD (A) and SLEs (B). All measurements used field potential recordings. **(A)** The onset of an SD was marked by beginning of a train of bursts that was followed by a sudden, large depolarization in whole cell recording or a large negative deflection in the field recording. The amplitude of an SD was measured as the maximal negative deflection relative to baseline. The half duration of SD was from the onset of the SD to the point where the recovery of SD reached half of its maximal amplitude. **(B)** The onset of an SLE was the beginning of the sudden large depolarization in whole cell recording, which corresponded to the initial burst in the field recording. The duration was the difference between the onset and the point where the SLE returned to baseline. The amplitude was the maximal negative deflection during the high-frequency bursts, which corresponded to the sustained firing in the whole cell recording.

**Supplementary Figure 2.**
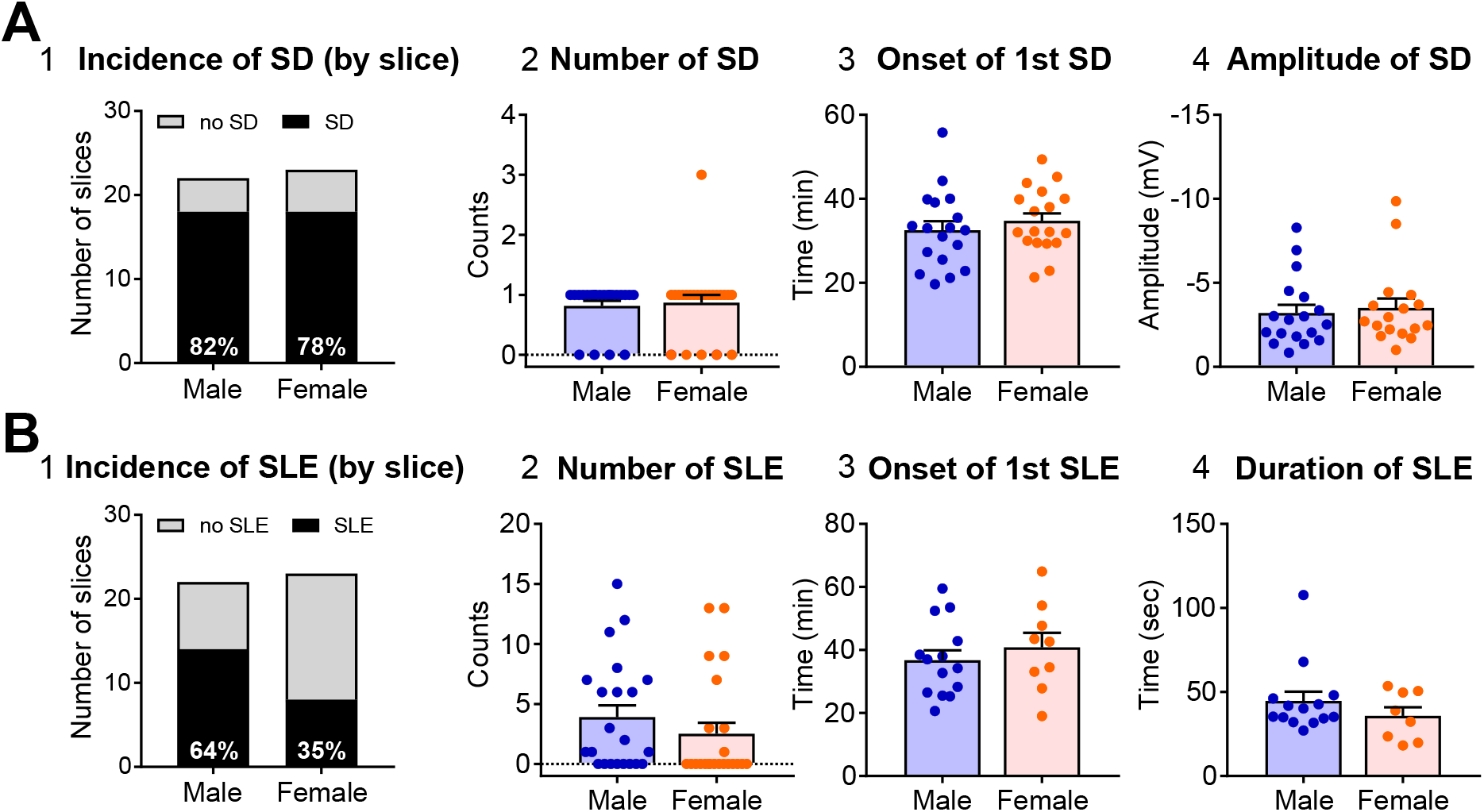
Lack of sex difference in characteristics of SD and SLEs in the first 60-min of 0 Mg^2+^/5 K^+^ aCSF exposure. **(A)** Quantification of SD. No statistical difference was found between males and females in the incidence of SD (A1, male, 22 slices/12 mice; female, 23 slices/10 mice; Fisher’s exact test, p > 0.9), number of SDs (A2, male, 22 slices/12 mice; female, 23 slices/10 mice; Mann-Whitney test, U = 253, p > 0.9), onset of the 1^st^ SD (A3, male, 18 slices/10 mice; female, 18 slices/9 mice; Unpaired t test, t(34) = 0.80, p = 0.43), and amplitude of SDs (A4, male, 18 slices/10 mice; female, 18 slices/9 mice; Mann Whitney test, U = 137, p = 0.61). **(B)** Quantification of SLEs. No statistical difference was found between males and females in the incidence (B1, male, 22 slices/12 mice; female, 23 slices/10 mice; Fisher’s exact test, p = 0.08), the number (B2, male, 22 slices/12 mice; female, 23 slices/10 mice; Mann-Whitney test, U = 191.5, p = 0.14), the onset (B3, male, 14 slices/9 mice; female, 8 slices/5 mice; Unpaired t test, t(20) = 0.20, p = 0.85), and the duration of SLEs (B4, 14 slices/9 mice; female, 8 slices/5 mice, Mann-Whitney test, U = 46, p = 0.53).

**Supplementary Figure 3.**
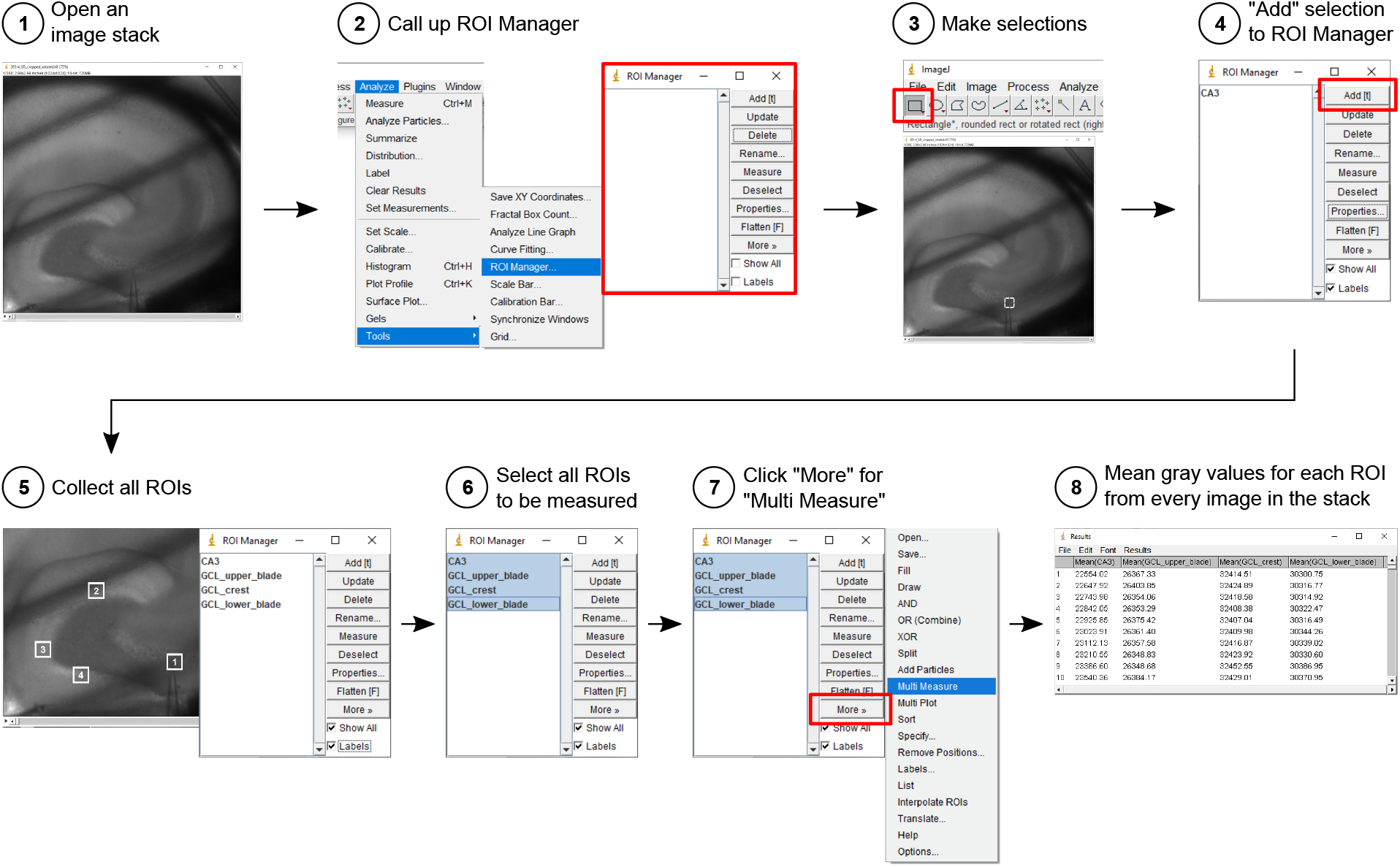
Intrinsic optical signal analysis using ImageJ. **Step 1:** Images acquired throughout an SD were processed as an image stack. **Step 2:** Region of interest (ROI) manager was used to organize multiple ROIs. **Step 3:** ROIs were 50 pixels x 50 pixels squares, drawn by the rectangle tool. **Step 4:** Each ROI was added into ROI Manager. **Step 5:** CA3 (#1), upper blade of granule cell layer (GCL) (#2), crest of GCL (#3), and lower blade of GCL (#4) were analyzed. **Step 6:** Once all ROIs were collected, all ROIs were selected for analysis all at one time. **Step 7:** Analysis was made using the “Multi Measure” function. **Step 8:** Mean gray values of each ROI from every image in the stack were measured and displayed in the result window. The list of ROIs was saved and re-used with proper adjustment for anatomical locations for the next image stack

## Supplementary Video

A recorded SD event is shown in a four-time faster fashion. Left: original intrinsic optical recording. Right: difference from baseline with pseudocolor. The timelapse images were recorded every 0.5 sec and replayed at the four-time of the speed. The propagation of SD wavefront is evident from CA3 to CA1 and to dentate gyrus.

